# Genome data artifacts and functional studies of deletion repair in the BA.1 SARS-CoV-2 spike protein

**DOI:** 10.1101/2024.01.23.575696

**Authors:** Miguel Álvarez-Herrera, Paula Ruiz-Rodriguez, Beatriz Navarro-Domínguez, Joao Zulaica, Brayan Grau, María Alma Bracho, Manuel Guerreiro, Cristóbal Aguilar-Gallardo, Fernando González-Candelas, Iñaki Comas, Ron Geller, Mireia Coscollá

## Abstract

Mutations within the N-terminal domain (NTD) of the spike (S) protein are critical for the emergence of successful SARS-CoV-2 viral lineages. The NTD has been repeatedly impacted by deletions, often exhibiting complex and dynamic patterns, such as the recurrent emergence and disappearance of deletions in dominant variants. This study investigates the influence of repair of NTD lineage-defining deletions found in the BA.1 lineage (Omicron variant) on viral success. We performed comparative genomic analyses of more than 10 million SARS-CoV-2 genomes from GISAID to decipher the transmission success of viruses lacking S:ΔH69/V70, S:ΔV143/Y145, or both. These findings were contrasted against a screening of publicly available raw sequencing data, revealing substantial discrepancies between data repositories, suggesting that spurious deletion repair observations in GISAID may result from systematic artifacts. Specifically, deletion repair events were approximately an order of magnitude less frequent in the read-run survey. Our results suggest that deletion repair events are rare, isolated events with limited direct influence on SARS-CoV-2 evolution or transmission. Nevertheless, such events could facilitate the emergence of fitness-enhancing mutations. To explore potential drivers of NTD deletion repair patterns, we characterized the viral phenotype of such markers in a surrogate *in vitro* system. Repair of the S:ΔH69/V70 deletion reduced viral infectivity, while simultaneous repair with S:ΔV143/Y145 led to lower fusogenicity. In contrast, individual S:ΔV143/Y145 repair enhanced both fusogenicity and susceptibility to neutralization by sera from vaccinated individuals. This work underscores the complex genotype-phenotype landscape of the spike NTD in SARS-CoV-2, which impacts viral biology, transmission efficiency, and immune escape potential, offering insights with direct relevance to public health, viral surveillance, and the adaptive mechanisms driving emerging variants.

## Introduction

As we navigate the constantly shifting landscape of the COVID-19 pandemic, the remarkable potential of SARS-CoV-2 for genetic adaptation has taken centre stage. Global turnover of SARS-CoV-2 lineages happened several times, with three variants of concern (VOCs) displacing the previous predominant lineages just in 2021: first, Alpha (B.1.1.7 and Q sublineages), then Delta (B.1.617.2 and AY sublineages), and lastly, Omicron (B.1.1.529 and BA sublineages, among others) (Rambaut et al. 2020). This successive replacement of lineages throughout the pandemic suggests that the newer had a higher adaptive value than the previous ones (Z. Chen et al. 2021; da Silva Francisco Junior et al. 2021), and thus, these three lineages are supposed to carry mutations associated with higher transmissibility or immune evasion.

The spike (S) protein —a type I transmembrane N-linked glycosylated protein— is a hotspot for mutations with high adaptive value. Spike proteins are located on the surface of SARS-CoV-2, and their main role is to mediate viral cell entry (Letko et al. 2020). The spike protein forms a homotrimer, which is cleaved post-transcriptionally into two subunits: S1 and S2. The S1 consists of the amino or N-terminal domain (NTD) and the receptor-binding domain (RBD), and it is responsible for binding to the host cell-surface angiotensin converting enzyme-2 (ACE2) receptors. The S2 subunit includes the trimeric core of the protein and is responsible for membrane fusion (Harvey et al. 2021; V’kovski et al. 2021). Therefore, some amino acid changes in the S protein may confer an advantage for transmission, considering its role in mediating viral cell entry. Additionally, most antibodies target sites either at the NTD or RBD and, therefore, mutations in these regions may enhance immune escape.

Specifically, deletions may play a decisive role in SARS-CoV-2 adaptive evolution, particularly in deletion-tolerant genome regions such as the S gene, as they can hardly be corrected by the proofreading activity of its RNA-dependent RNA polymerase (RdRP) (Denison et al. 2011; McCarthy et al. 2021; Minskaia et al. 2006). Notably, the NTD has been extensively impacted by deletions, which have emerged independently in multiple variants, often exhibiting complex and dynamic patterns, exemplified by the recurrent emergence and disappearance of the S:ΔH69/V70 deletion in dominant variants (McMillen et al. 2022; Meng et al. 2021). Acquisition of deletions in the NTD of the spike glycoprotein during long-term infections of immunocompromised patients has been reported and identified as an evolutionary pattern defined by recurrent deletion patterns that alter defined antibody epitopes (Avanzato et al. 2020; Choi et al. 2020). For example, different deletions are observed in Delta (S:ΔE156/F157-R158G), Alpha (S:ΔY144) and BA.1 (S:ΔV143/Y145) variants, all mapping to the same surface area, indicating a convergent function (Mishra et al. 2022; Planas et al. 2021). All these mutations are within the NTD site targeted by many anti-NTD neutralizing antibodies (McCallum, De Marco, et al. 2021). In the Alpha variant, S:ΔH69/V70 and S:ΔY144 are directly associated with antibody escape (Kemp et al. 2021; McCallum, De Marco, et al. 2021) and increased infectivity (Cantoni et al. 2022). More recent instances exist as well. Despite S:ΔY144 not being found in BA.2, its descendant lineage BJ.1 seems to have independently acquired this deletion (see cov-lineages/pango-designation issues #915 and #922 on GitHub). Then, it was passed on to XBB viruses through recombination with a BA.2-descended lineage, which was associated with increased immune escape regardless of the vaccination status or infection history (Mykytyn et al. 2023; Tamura et al. 2023; Zhang et al. 2022). Subsequently, in January 2023, the newly emerging XBB.1.5 lineage was shown to display an increased receptor-binding affinity and infectivity with respect to its parental lineage (Uriu et al. 2023; Yue et al. 2023).

The BA.1 lineage exhibited a growth advantage over previous variants of concern (Puhach et al. 2022; Ward et al. 2022) and soon took over the Delta variant. During this time, we became intrigued by the lack of overlap in NTD mutations between them. This is not the case for the Alpha variant and the BA.1 lineage, which share NTD deletions. If mutations in the NTD increased immune escape without compromising binding to ACE2, one might expect mutations in NTD to have a cumulative (“the more the better”) effect. However, that would not be the case if epistatic interactions between sites prevented the fixation of particular mutations in different genomic and genetic backgrounds. Our primary objective in this study was to investigate the effect of NTD deletion repair in BA.1 background. We examined the differences in success among three BA.1 haplotypes lacking the S:ΔH69/V70 and S:ΔV143/Y145 deletions, individually and in combination, which had been recurrently identified in immunocompromised individuals with chronic infections before widespread vaccination against COVID-19 (Avanzato et al. 2020; Choi et al. 2020; Kemp et al. 2021; McCarthy et al. 2021). Our approach integrates public data surveys with functional evidence to better understand how rare mutational events might shape viral evolution and pandemic dynamics.

## Materials and Methods

### GISAID survey data retrieval and processing

We performed two surveys to track the absence of NTD deletions S:ΔH69/V70 and S:ΔV143/Y145 in sequences assigned to the Omicron lineage BA.1 or any of its sublineages (hereafter referred to collectively as “BA.1”). The first survey included 11,334,504 available SARS-CoV-2 sequences fetched from the GISAID (Khare et al. 2021) EpiCoV database on June 15, 2022. The GISAID EPI_SET is available at DOI: 10.55876/gis8.230801ex. We ran Nextalign CLI v1.9.0 (Hadfield et al. 2018) with default scoring settings to obtain aligned genomes and peptides corresponding to each gene. Genomes from non-human hosts were discarded. Then, we filtered out genomes containing more than 5 % ambiguous bases and 1,000 gaps, or at least one indeterminate position among the lineage-defining sites of the spike protein. 10,353,158 genomes passed these filters. Then, BA.1 genomes were selected and classified into one of three haplotypes, depending on their lack of S:ΔH69/V70 (RepΔ69/70), S:ΔV143/Y145 (RepΔ143/145), or both (RepBoth). After running Nextclade CLI v3.8.2 (Aksamentov et al. 2021), we applied a BA.1 lineage filter alongside specific mutation and motif inclusion and exclusion criteria. Manual curation followed. For RepΔ69/70, sequences were selected if they carried S:ΔV143, S:ΔY144, either S:ΔG142 or S:ΔY145, and a lack of S:ΔH69 and S:ΔV70. Sequences were required to match the S:69-70:HV motif. For RepΔ143/145, sequences were selected if they carried S:ΔH69, S:ΔV70, either S:G142D or S:Y145D, and a lack of S:ΔV143, S:ΔY144, and either S:ΔG142 or S:ΔY145. The required motif was S:142-145:DVYY. For RepBoth, sequences were selected if they carried S:G142D, and a lack of S:ΔH69, S:ΔV70, S:ΔG142, S:ΔV143, S:ΔY144 and S:ΔY145. Sequences were required to match both the S:69-70:HV and S:142-145:DVYY motifs. We introduced an additional quality control step based on known control mutations: because BA.1 viruses typically exhibit the ORF1a:ΔS3675/F3677 (NSP6) deletion, sequences lacking this deletion were discarded. We used R v4.3.3 along with *tidyverse* v2.0.0 (Wickham et al. 2019) to conduct, manage and visualize these analyses.

To assess potential artifacts in the quality of GISAID data, we performed an initial exploratory analysis of raw sequencing reads obtained via the European Nucleotide Archive (ENA). First, we identified GISAID genomes with corresponding raw reads deposited in the ENA by cross-referencing GISAID identifiers (virus name and accession number). Raw reads were downloaded and reprocessed to test the congruence between the GISAID consensus sequences and the raw read mappings. The mappings were manually inspected to confirm genotypes, and we calculated two metrics for each haplotype: the ratio of fixed deletion repairs (allele frequency >0.95) and the ratio of cases with significant deletion repair read support (allele frequency >0.10). Both the exploratory analysis and full ENA survey workflow described hereunder employed the same workflow outline and software.

### ENA survey data retrieval and processing

Following this exploratory comparison, we extended the analysis to independently contrast the GISAID survey. We performed a second survey of combinations of NTD repaired deletions using SARS-CoV-2 “read run” records via the ENA Portal API. This survey tracked the absence of deletions S:ΔH69/V70 and S:ΔV143/Y145 (separately and combined) in sequences assigned to lineage BA.1 (Omicron variant). We searched for runs from samples collected from 1 November 2021 to 1 August 2022, filtering by NCBI taxonomy code (2697049 for SARS-CoV-2) and host scientific name (*Homo sapiens*). Instrument platforms matching DNBseq, Element and capillary sequencing were discarded. We also discarded runs with an RNAseq library strategy, as well as transcriptomic, metagenomic and metatranscriptomic library sources. The survey brought out 1,541,892 matching records as of 10 December 2024 (Supplemental Table 1). After filtering for data URL availability, 1,250,257 records remained. The date of submission of these records ranged between 10 November 2021 and 1 December 2024. We did a near 3-fold reduction of the dataset size, keeping 464,859 records selected at random for further analysis (see also Supplemental Table 1). Then, we ran an in-house pipeline implemented as a Snakemake (Köster & Rahmann 2012) workflow to detect NTD repaired deletions. We ran Snakemake v8.25.3 with Python v3.12.7.

For each sample, the following steps were carried out. FASTQ files were downloaded using the FTP URLs found in the ENA metadata. Then, read preprocessing and quality control (QC) were performed using *fastp* v0.23.4 (S. Chen 2023) with default settings. Filtered reads were mapped to a complete genome sequence from a BA.1 SARS-CoV-2 isolate (GenBank accession number: OZ070629.1) with *minimap2* v2.28 (H. Li 2018, 2021), using the recommended presets according to the software documentation (*sr* for Illumina and Ion Torrent reads, *map-ont* for Oxford Nanopore reads, and *map-hifi* for PacBio reads). After the mapping step, pileups were calculated using *samtools* v1.20 (Danecek et al. 2021) and consensus genomes were obtained with *iVar* v1.4.3 (Grubaugh et al. 2019) with a minimum insertion frequency of 0.9, a minimum depth of 40 and otherwise default settings. These sequences were fed to *pangolin* v4.3 (O’Toole et al. 2021) to perform lineage assignment. The output was used to further filter the dataset, keeping records with passing *pangolin* QC and an assignment of BA.1 lineage or Omicron (BA.1-like) constellation. Then, variant calling was performed on the remaining samples with *iVar* v1.4.3 as well, with a minimum frequency threshold of 0.05, a minimum depth of 40 and otherwise default settings. Results were further annotated using *SnpEff* v5.2 and filtered with *SnpSift* v5.2 (Cingolani et al. 2012). We used a custom *SnpEff* database based on the mapping reference. Finally, matching records were classified into one of the deletion repair haplotypes —RepΔ69/70, RepΔ143/145 or RepBoth— while records with no evidence of repaired deletions were discarded. We established a minimum allele frequency threshold of 0.75 for each repaired deletion allele. For the two single repair haplotypes, we set a maximum frequency threshold of 0.25 non-repaired deletion.

Additionally, we generated graphic visualizations of the genome region between sites 21694 and 22010 for visual inspection using R v4.3.3 with *ape* v5.8 (Paradis & Schliep 2019), *Rsamtools* v2.18.0 (Morgan et al. 2023), *tidyverse* v2.0.0 (Wickham et al. 2019) and *ggpubr* v0.6.0 (Kassambara 2023). The repository at https://github.com/PathoGenOmics-Lab/ena-spike-ntd-repdel-analysis (v1.0.0) contains the Snakemake workflow and the code to perform the initial screening.

### Number of emergences and transmission events

To infer the success of each combinatorial haplotype, we assessed the number of minimum emergence events and whether these originated subsequent transmission events. We placed genomes matching deletion repair haplotypes on a global-scale mutation-annotated tree (MAT) of public sequences (McBroome et al. 2021), under a maximum parsimony criterion, using UShER v0.6.2 (Turakhia et al. 2021). For GISAID data, genomes were placed on a tree dated to the survey date (15 June 2022), while for ENA data, consensus sequences inferred through our pipeline were placed on a tree dated to the last input record (11 March 2022).

To quantify the transmission of each haplotype, we developed *tcfinder* v1.0.0, a lightweight and efficient tool that performs an exhaustive breadth-first search of the samples on a phylogeny. This phylogeny must be provided using the tabular *phylo4* structure, as defined in *phylobase* v0.8.10 (https://github.com/fmichonneau/phylobase). The tool is available in Bioconda (Grüning et al. 2018). We selected clusters containing at least 2 members and at least 90 % target samples. This requirement was reduced from 100 % to account for potential sequencing errors and ambiguous sample placements on the phylogenetic tree.

The minimum number of independent emergences for each haplotype was derived from the phylogenies, by adding up the number of transmission clusters to the number of samples that were not transmitted. To adequately compare the relative epidemiological success of the most prevalent haplotypes, we developed a method to estimate their transmission fitness within each cluster. This was calculated as the ratio between the size of each transmission cluster and the number of sequences that were collected between the first and last cases of the cluster, with a 7-day padding added to both ends of the time window to account for missing data and short-lived clusters. This approach effectively standardizes the size of each transmission cluster, correcting differences in surveillance efforts and viral prevalence at different geographic regions and time periods. In essence, it provides a specific rate of emergence and transmission success relative to the broader population background. Consequently, for clusters exhibiting cross-border transmission (i.e., involving samples from different countries), we further divided the cluster time window into country-specific sub-windows. Then, the denominator of our transmission fitness estimate was calculated as the sum of the number of records in the data repository for each country-specific time window. For GISAID data, the full survey was used as background. For ENA data, the 464,859 records selected at random were used. Regardless, there were no significant differences between the distribution of fitness estimates within each haplotype, whether using country-specific sub-windows (“slicing” method) or not (“adding” method) (Supplemental Figure 1). Results in the main text use the first method. Differences in the distribution of the estimated transmission fitness between haplotypes and host age were evaluated using Wilcoxon rank-sum tests. To determine age fold changes, the ratio of group average values was calculated. Statistical analyses were performed and visualized using R v4.1.2 along with *tidyverse* v2.0.0 (Wickham et al. 2019) and *ggpubr* v0.4.0 (Kassambara 2023).

The complete pipeline (including the phylogenetic placement, transmission quantification, fitness estimation and data visualizations) is available as a Snakemake workflow at https://github.com/PathoGenOmics-Lab/transcluster (v2.1.0). We also studied the location and time span of the transmission clusters using the associated sample metadata from GISAID.

### Biological characterization of BA.1 deletion repair

Deletion repairs of S:ΔH69/V70 and S:ΔV143/ΔY145 were introduced individually and in combination into a pCG1 plasmid encoding a codon-optimized BA.1 spike protein (Giménez et al. 2022) by site-directed mutagenesis. All the constructs were verified by Sanger sequencing. Pseudotyped vesicular stomatitis virus (VSV) encoding a GFP reporter gene and carrying the different spike proteins was produced as previously reported (Gozalbo-Rovira et al. 2020). To assess the effects on virus production, pseudotyped VSV carrying each construct were produced independently three times. The resulting viruses were then titrated by infecting Vero E6 cells (kindly provided by Dr. Luis Enjuanes; CNB-CSIC, Spain) or Vero E6-TMPRSS2 cells (JCRB Cell bank catalogue code: JCRB1819) for 16 hours, followed by quantification of GFP-expressing infected cells using a live cell microscope (Incucyte SX5; Sartorius) to obtain the number of focus forming units (FFU) per millilitre. To assess thermal stability, 500 FFU of these pseudotyped viruses were incubated for 15 min at a range of temperature in a thermal cycler (30.4, 31.4, 33.0, 35.2, 38.2, 44.8, 47.0, 48.6 and 49.6 °C; Biometra T one Gradient, Analytik Jena) and the surviving virus was used to infect VeroE6-TMPRSS2 cells for 16 h. The GFP signal in each well was then determined using a live-cell microscope (Incucyte SX5, Sartorius). The average GFP signal observed in mock-infected wells was subtracted from all infected wells, followed by standardization of the GFP signal to the average GFP signal from wells incubated at 30.4 °C. Finally, we fitted a three-parameter log-logistic function to the data using the *drc* v3.0-1 R package (*LL.3* function) and calculated the temperature resulting in 50 % reduction in virus infection (*ED* function). To assess the effects on neutralization by polyclonal sera, we used six sera from convalescent patients from the first COVID-19 wave in Spain and six sera from individuals that had been administered two doses of the BioNTech-Pfizer Comirnaty COVID-19 vaccine. The neutralization capacity of the sera (NT50) was obtained as previously described on VeroE6-TMPRSS2 cells (Giménez et al. 2022). We used a previously described flow cytometry assay based on the use of polyclonal sera to examine surface expression (Grzelak et al. 2020). Briefly, HEK293T cells were transfected with the different S mutants using the calcium chloride method. 24 h later, cells were detached using PBS with 1 mM EDTA, washed, and incubated in ice with different polyclonal sera (three from convalescent patients from the 1st COVID-19 wave in Spain and one from individuals that had been administered two doses of the BioNTech-Pfizer Comirnaty COVID-19 vaccine) at a 1:300 dilution in PBS containing 0.5 % BSA and 2 mM EDTA for 30 min. Next, cells were washed three times with PBS, stained with anti-IgG Alexa Fluor 647 (Thermo Fisher Scientific) at a 1:400 dilution, and analyzed by flow cytometry similarly treated un-transfected controls to set the threshold for positive cells. For cell-cell fusion assays we used a split Venus fluorescent protein system (García-Murria et al. 2019). HEK293T cells were grown overnight in 24 well plates (1.5 x 105 cells/well) using DMEM supplemented with 10 % FBS. After 24 hours, cells were transfected using Lipofectamine 2000 (Invitrogen) with 0.5 µg of either a 1:1 mixture of the S plasmids and a Jun-Nt Venus fragment (Addgene 22012) plasmid or a mixture of hACE2 plasmid (kindly provided by Dr. Markus Hoffman; German Primate Center, Goettingen/Germany) (Hoffmann et al. 2020) and the Fos-Ct Venus fragment (Addgene 22013). After 24 h, cells were counted, and the S-transfected cells were mixed at a 1:1 ratio with ACE2-transfected cells and seeded in 96 well plates (3 x 104 cells/well) in 100 µL of media. Cells transfected with the Wuhan-Hu-1 served as a positive control, while cells transfected with hACE2 and Jun-Nt Venus were used as negative control. We obtained the GFP Integrated Intensity (GCU·µm²/image) in each condition using a live-cell imaging platform (Incucyte SX5, Sartorius) at 24 hours post mixing and standardized to the signal obtained from the positive control (Wuhan-Hu-1 spike protein). All experiments were performed at least three times in triplicates.

Statistical analyses were performed and visualized using R v4.1.2, along with *tidyverse* v2.0.0 (Wickham et al. 2019) and *ggpubr* v0.4.0 (Kassambara 2023) to facilitate the analysis and enhance the visualization of these results. Comparisons were conducted utilizing t-tests. We used unpaired tests for all assays but neutralization (log-transformed NT50 values) and surface expression, for which we used paired tests, as we used the same sets of polyclonal sera in each experiment. To determine fold change values, the ratio of group average values was calculated.

## Results

### Detection and transmission dynamics of spike NTD deletion repair in the BA.1 lineage using GISAID genomes

We first examined the differences in success within the Omicron BA.1 lineage depending on the presence of ΔH69/V70 and ΔV143/Y145 in the NTD of the spike protein through a global survey of viral genome data. This domain is characterized by recurrent deletions occurring in distinct independent lineages (McCallum, De Marco, et al. 2021; McCallum, Walls, et al. 2021; McCarthy et al. 2021; Meng et al. 2021; Planas et al. 2021; Tamura et al. 2023). We interrogated the possibility of different patterns of deletion repair of epidemiological significance and the drivers behind their emergence through a survey of 11.3 million samples from the GISAID’s EpiCov platform (Figure 1A). We identified 5,499 samples that carried repaired S:ΔH69/V70, S:ΔV143/Y145, or both (Figure 2A, Table 1; full details in Supplemental Table 2). The most frequently observed group consisted of samples exhibiting the repaired S:ΔH69/V70 in BA.1 background, while the remaining repair patterns were less frequently observed.

**Figure 1.**
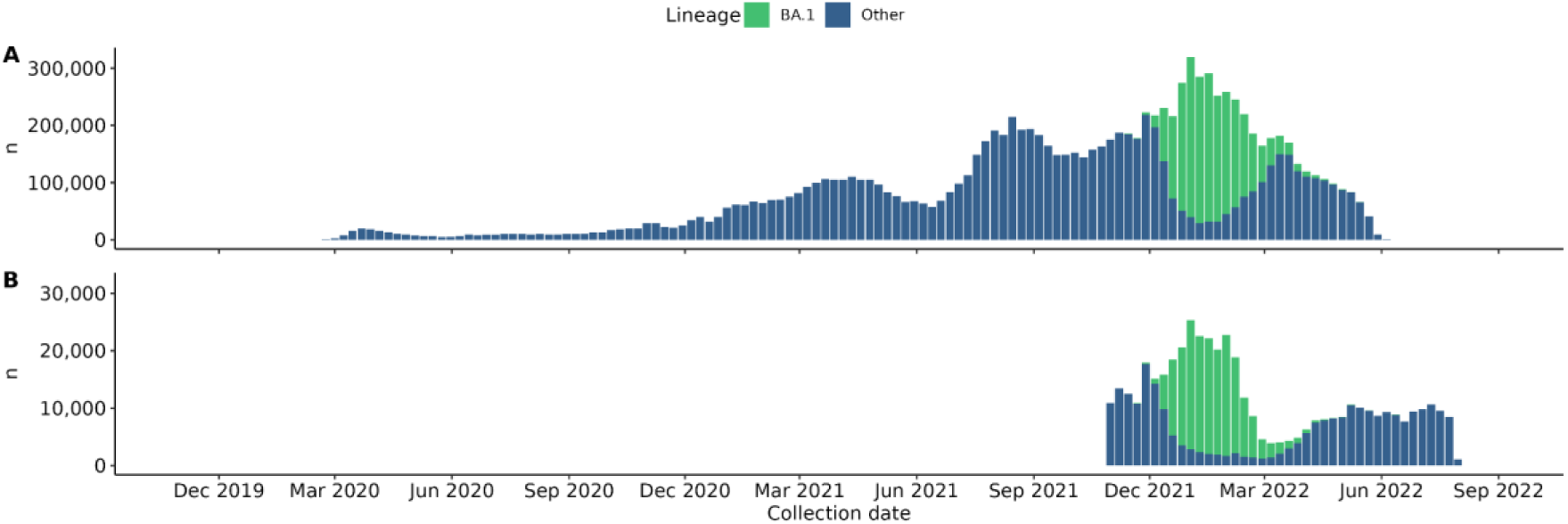
Distribution of collection dates of the two survey datasets. Bars show the weekly number of records in each survey dataset, excluding those with missing collection dates. Records identified as deletion repair-bearing were extracted from the BA.1 data partition in both surveys, which is highlighted in a lighter shade. (A) GISAID dataset, with lineages extracted from metadata. (B) ENA dataset (3-fold random subset), with lineages assigned running *pangolin* v4.3 on consensus sequences reconstructed from read mappings (see Materials and Methods), as no lineage metadata is provided in database records. Although lineage filtering prior to data processing was not possible for the ENA dataset, the chosen time frame ensured adequate representation of BA.1 records.

**Figure 2.**
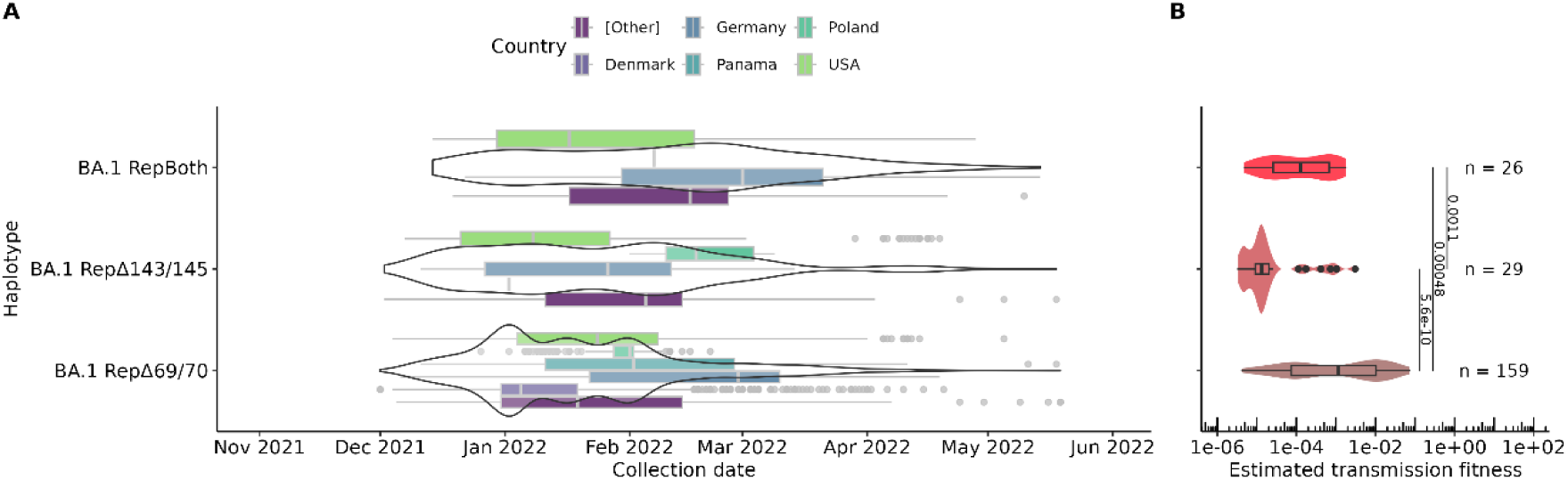
Summary of GISAID survey results for BA.1 genomes with repaired S:ΔH69/V70, S:ΔV143/Y145 or both. A) Sample collection timelines broken down by sampling location. The five countries with the highest number of observations globally are displayed. B) Differences in the estimated transmission fitness between deletion repair haplotypes. This metric corrects the size of each transmission cluster regarding local spatial and temporal sequencing efforts and lineage abundances. The differences were assessed using Wilcoxon rank-sum tests, and P-values are displayed above each line connecting a pair of haplotypes. Each data point corresponds to a transmission cluster. Group sizes (number of transmission clusters, n) are displayed next to the p-values.

**Table 1.**
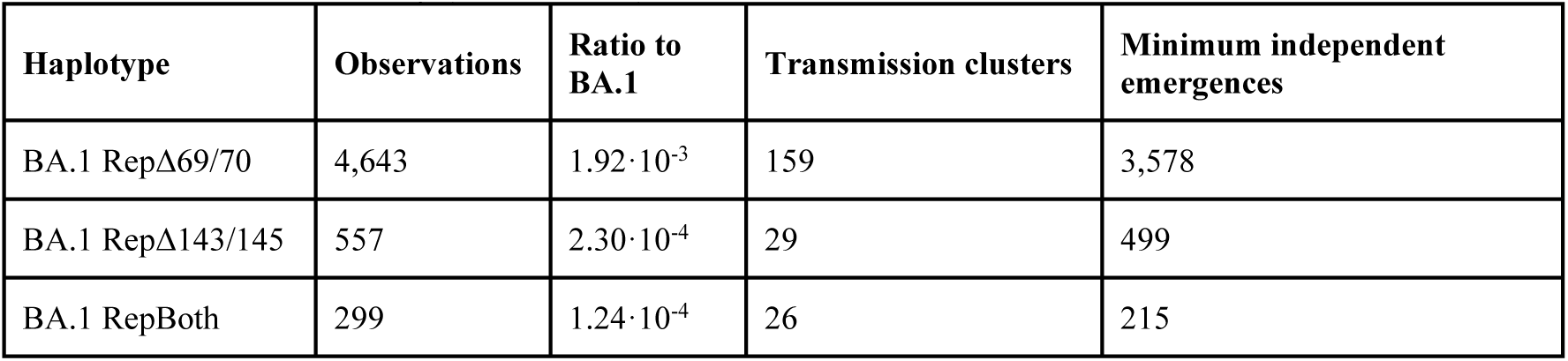
Description of Omicron BA.1 haplotypes with repaired NTD deletions detected through the GISAID survey. The ratio to BA.1 observations is calculated as the fraction of observations over the number of BA.1 observations found in the survey (n = 2,420,866).

Nevertheless, estimates of prevalence can be distorted by several factors, including the geographical location and temporal specificity associated with both sequencing efforts and the distribution of lineages throughout the pandemic. Transmission, on the other hand, is usually considered as a proxy of viral fitness, because it is related to its basic reproductive number, reflecting the ability of the virus to replicate, persist and spread within hosts and in the population (Domingo 2016). Therefore, we identified the transmission clusters of each haplotype through a phylogenetic estimation using worldwide SARS-CoV-2 genomes. We measured an average within-cluster collection date window of 24 ± 27 days, with 70 % of clusters showing only within-border transmission (Supplemental Figure 2). The haplotype bearing the S:ΔH69/V70 single repair had the highest median transmission fitness, followed by the double repair and, finally, by the S:ΔV143/Y145 single repair (Figure 2B). We then investigated host age as a potential adaptive determinant of NTD deletion repair in BA.1 background. No significant differences in host age were observed between deletion repair haplotypes (Supplemental Figure 3). However, transmitted samples with repaired S:ΔH69/V70 were associated with higher host age compared to non-transmitted samples (P = 4.7·10^⁻3^; Supplemental Figure 4), an effect likely independent of sampling location (Supplemental Figures 5 and 6). These findings should be interpreted cautiously due to substantial missing patient metadata (Supplemental Figure 7).

To evaluate the reliability of GISAID-derived data, we compared its consensus sequences to corresponding raw reads from the European Nucleotide Archive (ENA). We cross-referenced the available ENA entries with all GISAID identifiers of genomes with deletion repairs to identify matching records (Supplemental Table 3). Among 18 genomes with matching ENA runs, less than 25 % confirmed fixed deletion repairs as indicated by the GISAID consensus, and fewer than 60 % had significant read support of the deletion repair genotype. Specifically, for repaired S:ΔH69/V70, seven genomes matched ENA runs, with only one exhibiting a fixed repaired deletion. For repaired S:ΔV143/Y145, one genome matched ENA runs, with none showing fixed repairs. For genomes with both repaired deletions, ten matched ENA runs, with three displaying fixed repairs. These findings suggest that GISAID sequences may not reliably represent deletion repair genotypes, highlighting the need for further validation of such analyses.

### Detection and transmission dynamics of spike NTD deletion repair in the BA.1 lineage using raw sequencing data

In our survey of viral genome data from GISAID we were able to identify repairs to deletion mutations, which are theoretically rare events. However, their validity may be confounded by sequencing artifacts. Public consensus genome sequences often suffer from low coverage, poor sequencing quality, and imputed data, hindering the detection of genuine deletion repair events. To address potential artifacts, we analyzed 464,859 read run records from the ENA (Figure 1B), selected out of 1.54 million collected between 1 November 2021 and 1 August 2022. This allowed us to reassess haplotype assignments with greater reliability.

We observed substantial discrepancies compared to the initial GISAID genome survey. We identified 24 samples that carried repaired S:ΔH69/V70, S:ΔV143/Y145 or both (Figure 3A, Table 2; full details in Supplemental Table 4) with an average haplotype-defining allele frequency of 0.918 (see Figure 3B). The ratio of repaired deletions relative to BA.1 observations was around 3- to 51-fold lower (depending on the haplotype) than in the GISAID dataset, underscoring a significant drop in detection. This suggests that the earlier survey was influenced by artifacts or biases in sequence processing.

**Figure 3.**
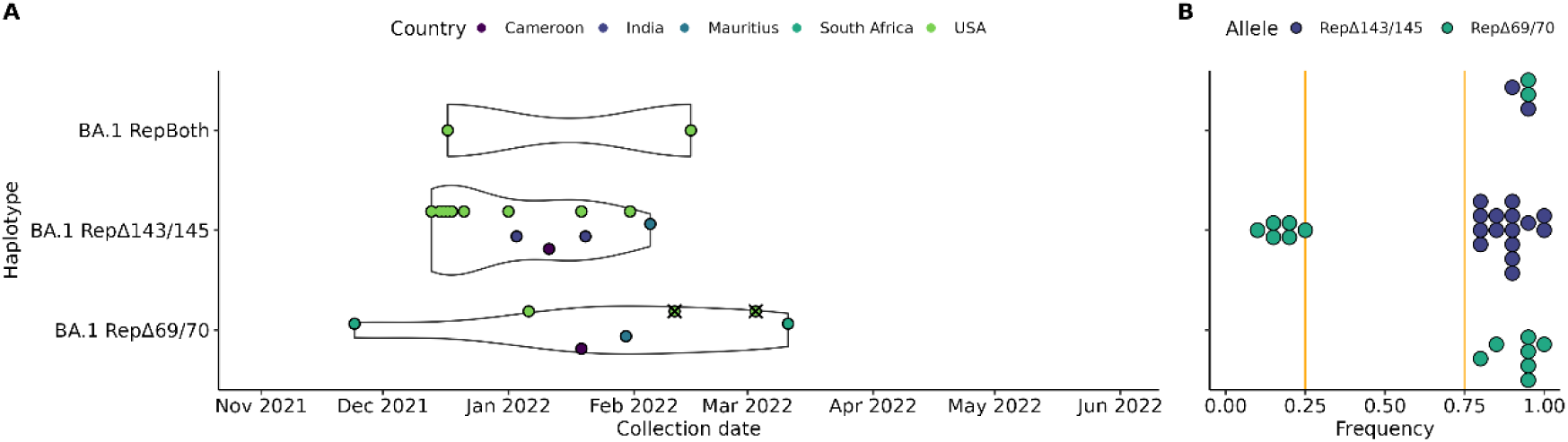
Summary of ENA survey results for BA.1 records with repaired S:ΔH69/V70, S:ΔV143/Y145 or both. A) Sample collection timelines broken down by sampling location. The five countries with at least one observation are displayed. Crossed markers indicate observations belonging to the single detected transmission cluster. The timeframe aligns with Figure 2A for consistency. B) Allele frequency of deletion repairs in each run record. For each allele, each point represents the allele frequency in a different record. Point bins have a width of 0.05. Note that, for haplotype RepΔV143/Y145, some samples bear repaired S:ΔH69/V70 with a frequency under the 0.25 threshold (see Materials and Methods). No differences in the estimated transmission fitness between deletion repair haplotypes are reported, as we did not detect transmission for all haplotypes (see Table 2).

**Table 2.**
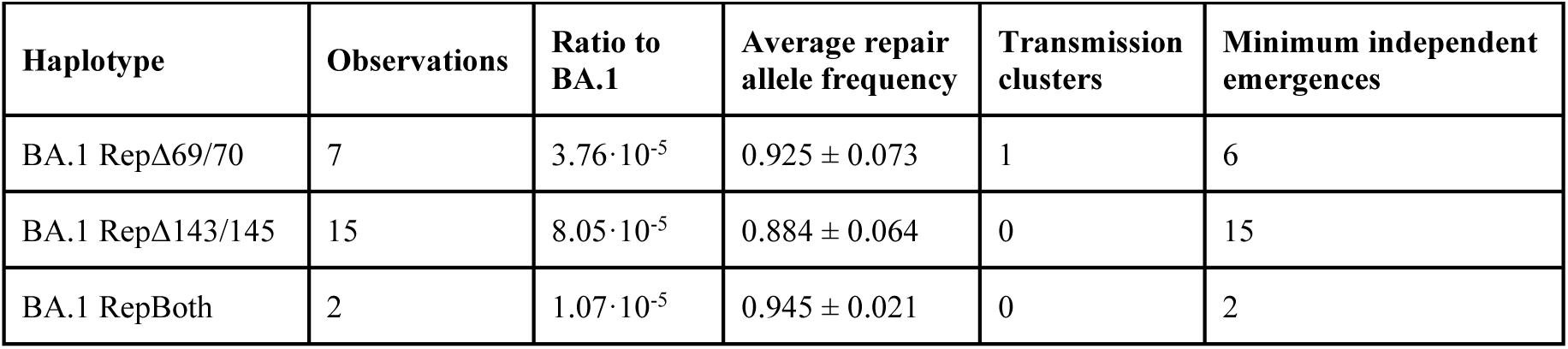
Description of Omicron BA.1 haplotypes with repaired NTD deletions detected through the ENA survey. The ratio to BA.1 observations is calculated as the fraction of observations over the number of BA.1 read runs found in the survey analysis (n = 186,387). The average and standard deviation (SD) of the frequency of the deletion repair allele(s) associated with each haplotype are reported as well.

We then performed a phylogenetic estimation of transmission clusters using worldwide SARS-CoV-2 genomes. A single transmission cluster was detected (see Table 2), involving two records with repaired S:ΔH69/V70 (accession numbers SRR19566289 and SRR20496144) collected 20 days apart in California and New Jersey, U.S. The allele frequency of the deletion repair declined 12 % between infections. Notably, the assigned haplotype exhibited the highest estimated transmission fitness in the earlier GISAID survey (see Figure 2B). In short, generally, each observation was detected as an independent emergence with limited epidemiological impact, suggesting that these NTD repaired deletion events might be rare, isolated, and unlikely to drive SARS-CoV-2 evolution or transmission dynamics by themselves.

### NTD deletion repair in Omicron BA.1 background has a non-accumulative effect on viral phenotype

NTD deletion repair events could have still contributed to viral diversity in a broader evolutionary context. Even isolated events might provide new opportunities for fitness-enhancing mutations to emerge, particularly in high-transmission scenarios. To investigate whether certain viral characteristics act as drivers of certain patterns of deletion repair compared to others, we analysed the effect of these events in the Omicron BA.1 spike using a pseudotyped VSV system. We generated VSV pseudotyped with the BA.1 spike proteins with repaired S:ΔH69/V70, repaired S:ΔV143/Y145, or both deletions repaired. We then used these pseudotyped viruses to examine possible phenotypic effects on spike function, including the efficiency of virus production, thermal stability, surface expression, susceptibility to antibody neutralization, and fusogenicity.

To assess possible effects on virus production, which encompasses both viral egress and infection of the next cell, VSV was pseudotyped with all spike constructs under identical conditions at the same time, and the amount of virus produced titrated on Vero E6 cells (Figure 4A). Virus production was unaffected by S:ΔV143/Y145 repair (0.77-fold change; P = 0.06) but was significantly reduced upon repairing S:ΔH69/V70 (0.28-fold; P = 6.6·10^-4^) or both deletions (0.34-fold; P = 1.6·10^-4^). A similar effect was observed when the titre of the virus was evaluated in cells expressing the TMPRSS2 co-receptor, indicating a substantial negative effect of S:ΔH69/V70 repair by itself on virus production (0.26-fold; P = 0.017).Reduced viral production can stem from lower stability of the Spike proteins or lower expression on the cell surface. To test the former, we obtained the temperature resulting in a 50 % reduction of virus titre for each spike variant (i.e., 50 % inactivation temperature; Figure 4B). No significant differences were observed, suggesting that spike protein stability was not the driver of these differences. We therefore examined whether the different constructs had altered cell expression of the different constructs by flow cytometry, using polyclonal sera from four individuals (Figure 4C). We did not detect differences in the median cell surface expression of the spike protein between the canonical BA.1 protein and any deletion repair haplotype. However, the repair of S:ΔV143/Y145 resulted in higher expression than S:ΔH69/V70 (3.2-fold; P = 0.0030) and the double repair (1.9-fold; P = 0.032), resembling the effect observed with virus production.

**Figure 4.**
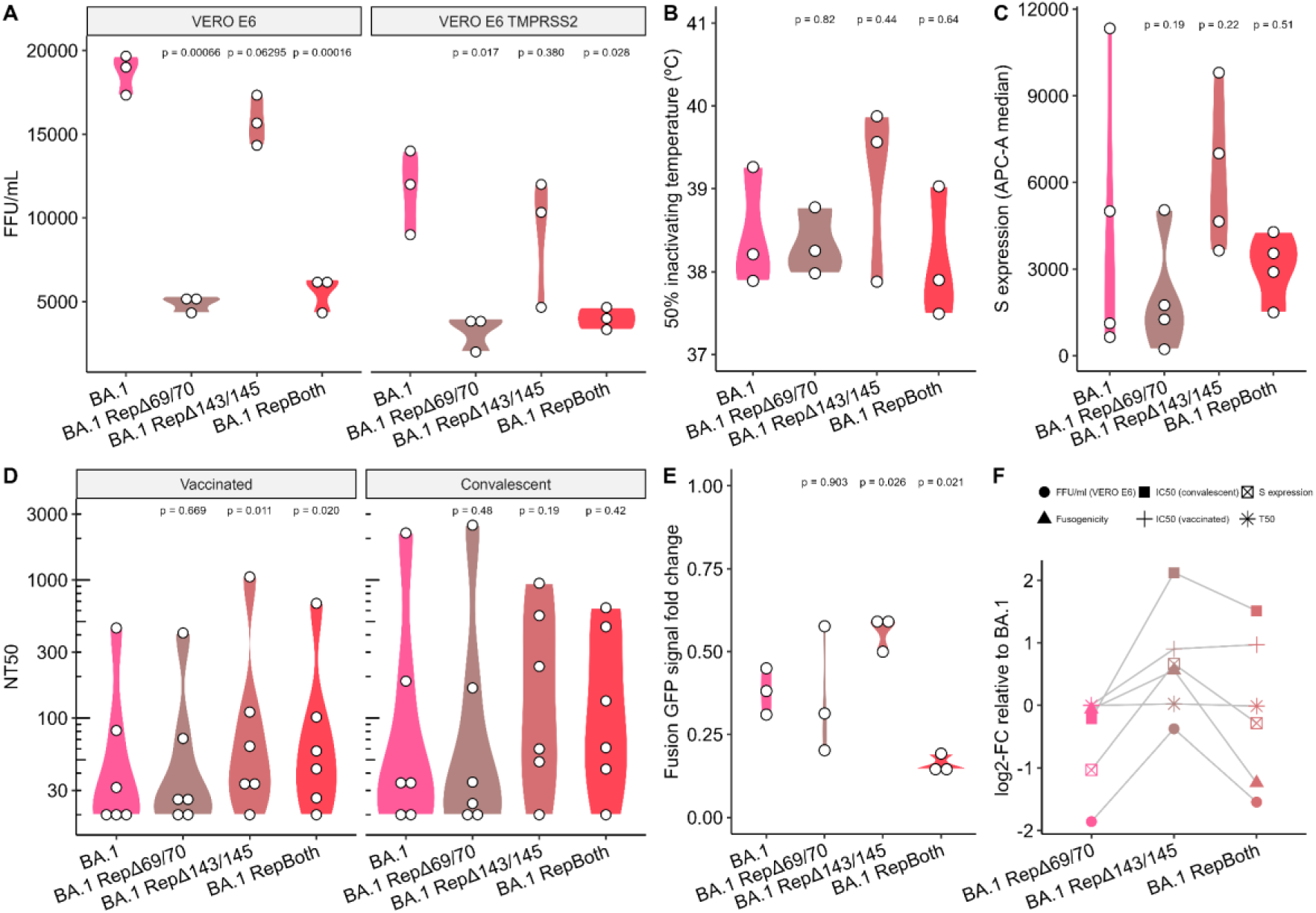
Experimental assessment of the effect of repairing NTD deletions in Omicron BA.1 background. (A) Comparison of viral titres obtained for each spike variant using pseudotyped VSV on the indicated cell line. (B) Comparison of the temperature resulting in 50 % inactivation of pseudotyped VSV carrying the indicated S protein. (C) Surface expression of the spike protein in transfected HEK293T cells quantified using flow cytometry. (D) Reciprocal 50 % neutralization titres (NT50) of sera from convalescent and vaccinated individuals. (E) Relative ability of each spike variant to drive cell-cell fusion relative to the Wuhan-Hu-1 spike. (F) Summary of the log2-transformed fold change for each spike variant relative to that of BA.1 in terms of average virus production in VERO E6 cells, average fusogenicity, median NT50, average surface expression and average 50 % inactivating temperature. Subfigures A-E represent the median and interquartile range of at least three replicates. Differences were assessed using Wilcoxon rank-sum tests.

As neutralizing antibodies can be key drivers of viral evolution, we questioned whether deletion repair affected neutralization by polyclonal sera from convalescent donors from the first epidemic wave in Spain and those dually vaccinated with the Comirnaty mRNA vaccine (n = 6 each; Figure 4D). Significant increases in susceptibility to neutralization in sera from vaccinated individuals against viruses with repaired S:ΔV143/Y145 (2.1-fold; P = 0.011) or both S:ΔH69/V70 and S:ΔV143/Y145 repaired (1.5-fold; P = 0.020) were observed. A similar trend was observed with convalescent sera, but statistical significance was not reached (P = 0.19 and P = 0.42, respectively) possibly due to a higher intra-sample variability. Thus, these results indicate that increased neutralization is driven by the repair of S:ΔV143/Y145 in BA.1 background.

Cell to cell spread via fusion of the plasma membrane could potentially reduce exposure to neutralizing antibodies, compensating for increased susceptibility to neutralization. Hence, we also examined whether deletion repair could alter the ability of the spike protein to fuse cells (Figure 4E). Interestingly, the repair of S:ΔV143/Y145 increased cell fusion relative to the BA.1 spike protein (1.5-fold; P = 0.026), while the repair of both S:ΔH69/V70 and S:ΔV143/Y145 led to a decrease of more than 50 % in the average fusogenicity (0.42-fold; P = 0.021). The presence of S:ΔH69/V70 by itself did not seem to play a role in cell-cell fusion in a BA.1 background.

## Discussion

Deletions in the SARS-CoV-2 genome have a significant impact on viral adaptation and fitness, often surpassing the effects of single nucleotide variants (Cantoni et al. 2022; McCarthy et al. 2021; Meng et al. 2021; Mishra et al. 2022; Venkatakrishnan et al. 2023). In fact, deletions in the NTD region of the spike protein are fixed in many prominent viral variants. Our understanding of the repair patterns of deletions is crucial for elucidating the genetic and phenotypic characteristics of the variants of concern. In this work, we demonstrate that repairing these deletions can alter viral characteristics and potentially influence the success, emergence and transmission dynamics of specific viral haplotypes.

The *in vitro* assessment of the viral characteristics driving different patterns of deletion repair in BA.1 showed differences in spike virus production and infectivity, fusogenicity, and sera neutralization, but no differences in thermal stability. Repair of S:ΔH69/V70 led to reduced virus titres individually and in combination with S:ΔV143/Y145 repair. We did not observe differences in thermal stability with the BA.1 protein, suggesting that the effect of S:ΔH69/V70 and S:ΔV143/Y145 repair did not affect full-protein stability.

Cell-cell fusion and subsequent formation of syncytia can be mediated by viral membrane glycoproteins (W. Li et al. 2003; Tseng et al. 2005) such as the spike protein found in SARS-CoV-2, and it is a key feature of SARS-CoV-2 infection (Buchrieser et al. 2020; Bussani et al. 2020; Hoffmann et al. 2020). In line with prior research, our results point to a lower fusogenicity of the unaltered BA.1 spike protein compared to that of Wuhan-Hu-1, which has been attributed to its inefficient spike cleavage and unfavoured TMPRSS2-mediated cell entry (Meng et al. 2021). Interestingly, deletion repair patterns exerted a non-accumulative influence on the fusogenicity of the BA.1 spike protein: repair of S:ΔH69/V70 alone did not have a significant impact and repair of S:ΔV143/Y145 promoted higher fusogenicity, while repair of both deletions led to a significant decrease in fusogenicity.

Deletion repair did also affect sera neutralization. In the BA.1 background, repaired S:ΔV143/Y145 was associated with a significant increase in sensitivity to neutralization by polyclonal sera obtained from vaccinated individuals. Conversely, we did not detect a significant association of repaired S:ΔV143/Y145 with sensitivity to sera from convalescent, “first-wave” patients. Repair of S:ΔH69/V70 did not affect neutralization with either serum group. Although our surrogate system might not fully reflect the complex interactions of the spike protein with host cells and the immune system *in vivo*, and other regions of the spike protein or the viral genome might also contribute to SARS-CoV-2 fitness and adaptation, our results point to the variant-independent influence of NTD variability on immune effects. These findings point to deletion repair events being rare and isolated, as their context-dependent and often deleterious effects generally suggest limited adaptive advantage. This may also indicate epistatic incompatibilities that constrain their contribution to viral fitness and transmission. In agreement with the *in vitro* results, sequence surveys suggest that deletion repair events were rare and likely isolated, with limited direct impact on SARS-CoV-2 evolution or transmission dynamics. This aligns with the hypothesis that recurrent deletion and apparent repair events in early variants of concern (VOC) may have been driven by persistent viral reservoirs, where prolonged infections or animal reservoirs allow for cycles of mutation accumulation and reversion before reintroduction. In contrast, such dynamics appear less relevant in second-generation dominant lineages (e.g., BA.2 sublineages), as reviewed by Roemer et al. (Roemer et al. 2023).

All survey efforts are expected to be biased by geographic and temporal differences in sequencing efforts and lineage prevalence. Sequence analysis could be affected by sampling bias and technical artifacts. However, despite our attempts to account for demographic differences between variants, our analysis could be affected by biases in genome sequencing and publication practices.

In our study, we initially observed instances of apparent genetic reversions that could, in fact, be attributable to missing information in poorly sequenced regions or polymorphic sites. The ENA survey revealed a significantly lower frequency of deletion repair events compared to GISAID, and the majority of one-to-one comparisons of GISAID genomes against raw reads data did not confirm the deletion repair genotype. This suggests potential artifacts in GISAID sequences, likely arising from a replacement of problematic genomic regions with the homologous ancestral sequence. Such undocumented adjustments can create reference bias, distorting the interpretation of genetic events and complicating genomic analyses, particularly given the quality control mechanisms employed by repositories like GISAID. In fact, key global resources such as the daily-updated mutation-annotated trees we used for transmission estimation (McBroome et al. 2021) could well be impacted by these issues. We highlight the need for greater scrutiny of sequence deposition and quality control mechanisms. Ongoing efforts such as the Viridian initiative (Hunt et al. 2024) have already pointed out these issues and are working toward reducing their impact. Incidentally, the limitations of the FASTA format itself also contribute to this problem, since it restricts the amount of metadata and quality information that can directly accompany sequences. This choice, while historically convenient, may no longer align with the need for richer data representations. Accordingly, submission of raw sequencing data from genome studies to public repositories should be prioritized.

We also acknowledge that our study does not specifically address the mutational mechanism that led to deletion repair. We speculate that recombination between different infecting variants could be behind the emergence deletion-repairing haplotypes, at least in some cases. We further speculate that template switching at wrong loci during replication (“imperfect homologous recombination”, as hypothesized by Chrisman et al. 2021) could explain the emergence of rare indel events such as deletion repairs in the absence of coinfections. Recombination might have played a role in the evolution of SARS-CoV-2 from the ancestral coronavirus before diversification in humans (Andersen et al. 2020; Boni et al. 2020; Kirtipal et al. 2020; MacLean et al. 2021; Wells et al. 2021), and studies using focused datasets have also pointed to ongoing recombination and proposed insights into its dynamics (e.g., Ignatieva et al. 2022; Jackson et al. 2021; Perez-Florido et al. 2023; VanInsberghe et al. 2021). Still, the emergence of the first demonstrably recombinant lineages of epidemiological relevance was not reported until years into the pandemic (Tamura et al. 2023; Yue et al. 2023). Due to the poor phylogenetic resolution of the datasets, our efforts ultimately led to an inconclusive result, rendering us unable to articulate a systematic verification of the recombination hypothesis.

In summary, our study describes the emergence of rare deletion repair events in the SARS-CoV-2 spike NTD, and investigates their impact on viral fitness, transmission, and clinical characteristics depending on the genetic context. In doing so, we have provided novel insights into the repair patterns of NTD deletions in Omicron BA.1 background and their phenotypic consequences. While significant progress has been recently made in estimating the fitness effects of individual mutations across the SARS-CoV-2 genome (Bloom & Neher 2023), these approaches do not focus on how mutations interact within different genomic contexts. There have also been efforts to specifically characterize the variability of the S protein NTD through full swap assays, comparing the ancestral (Wuhan-Hu-1) spike protein with that of Alpha and Omicron BA.1 (Cantoni et al. 2022), Delta and Omicron BA.1 and BA.2 (Meng et al. 2021) and even with more distant virus like SARS-CoV (Qing et al. 2021). However, to our best knowledge, no previous study has undertaken a combinatorial approach to characterize each separate marker. Our work reveals that repair of specific deletions might be driven by distinct phenotypic traits, such as changes in viral characteristics, including fusogenicity, infectivity and susceptibility to neutralization. Understanding these genotype-phenotype relationships can inform the evolutionary dynamics and adaptation of SARS-CoV-2 variants. Future studies building upon these findings will contribute to the development of effective strategies to monitor and mitigate the impact of NTD deletions in emerging SARS-CoV-2 variants.

## Data availability

The GISAID EPI_SET underlying this article is available via https://dx.doi.org/10.55876/gis8.230801ex. Raw reads data is publicly available via the European Nucleotide Archive (ENA) under the accession numbers provided in Supplemental Table 1. The software we developed and used to find clusters in a global phylogeny, *tcfinder* v1.0.0, is a command line tool released as free software under the GNU GPLv3 license. Its source code is available in GitHub (https://github.com/PathoGenOmics-Lab/tcfinder). It is also available as a package in the Bioconda *conda* channel (https://anaconda.org/bioconda/tcfinder). The pipeline used for the transmission analysis, *transcluster* v2.1.0, is a Snakemake workflow also released as free software under the GNU GPLv3 license, and its source code is available in GitHub (https://github.com/PathoGenOmics-Lab/transcluster). The processed data, code, configuration files and results of the ENA survey and both transmission analyses are available via https://dx.doi.org/10.20350/digitalCSIC/17032. The lists of sample identifiers assigned to each haplotype under study are also incorporated into the article and its online supplemental material. Code for performing the ENA survey and the pipeline for processing the resulting raw sequencing data is available as free software under the GNU GPLv3 license, and its source code is available in GitHub (https://github.com/PathoGenOmics-Lab/ena-spike-ntd-repdel-analysis; v1.0.0).

## Acknowledgements

We gratefully acknowledge all data contributors, i.e., the Authors and their Originating laboratories responsible for obtaining the specimens, and their Submitting laboratories for generating the genetic sequence and metadata and sharing via the GISAID Initiative, on which this research is based. We also extend our acknowledgement to all contributors and submitters of raw reads hosted by the European Nucleotide Archive (ENA) and other members of the International Nucleotide Sequence Database Collaboration (INSDC), which are also integral to this study. We thank Dr. Óscar González Recio (INIA-CSIC, Spain) for providing sequencing data from a BA.1 sample with S:142G and repaired NTD deletions (GISAID accession code: EPI_ISL_9805648) for an initial exploratory analysis related to S gene target failure (SGTF), Dr. Luis Enjuanes (CNB-CSIC, Spain) for providing Vero E6 cells, and Dr. Markus Hoffman (German Primate Center, Germany) for providing the hACE2 plasmid. We thank Dr. Irving Cancino Muñoz for his insightful feedback and recommendations on sequence data processing. We are grateful to Jimena Solana González for reviewing the manuscript and offering thoughtful feedback. We also thank Francisco José Martínez Martínez for comments on phylogeny visualization and analysis of transmission clusters. The computations were performed on the HPC cluster Garnatxa at the Institute for Integrative Systems Biology (I^2^SysBio, Spain). The I^2^SysBio is a joint collaborative research institute involving the University of Valencia (UV) and the Spanish National Research Council (CSIC).

## Ethics statement

Sera samples for the biological characterization of BA.1 deletion repair were obtained from the La Fe University and Polytechnic Hospital of Valencia and were collected after informed written consent had been obtained, with approval by the ethical committee and institutional review board (registration number 2020-123-1).

## Contributions

MAH, BND, PRR and MC conceived the theoretical framework. MAH and BND devised the initial idea. MAH and MC conceived the final study design. MAH implemented and performed the data retrieval, data curation and computations. JZ, BG, and RG designed the experiments. JZ and BG performed the experiments. MAH carried out the statistical analyses of the experiments. MG and CAG contributed biological samples. MAB did project management. MAH drafted the manuscript with support from BDN, PRR, RG and MC. FGC aided in interpreting the results. MAH, PRR, BND, RG, FGC, IC and MC discussed the results and commented on the manuscript. MC supervised the project.

