## Supplemental Figure 1 for "Genome data artifacts and functional studies of deletion repair in the BA.1 SARS-CoV-2 spike protein"

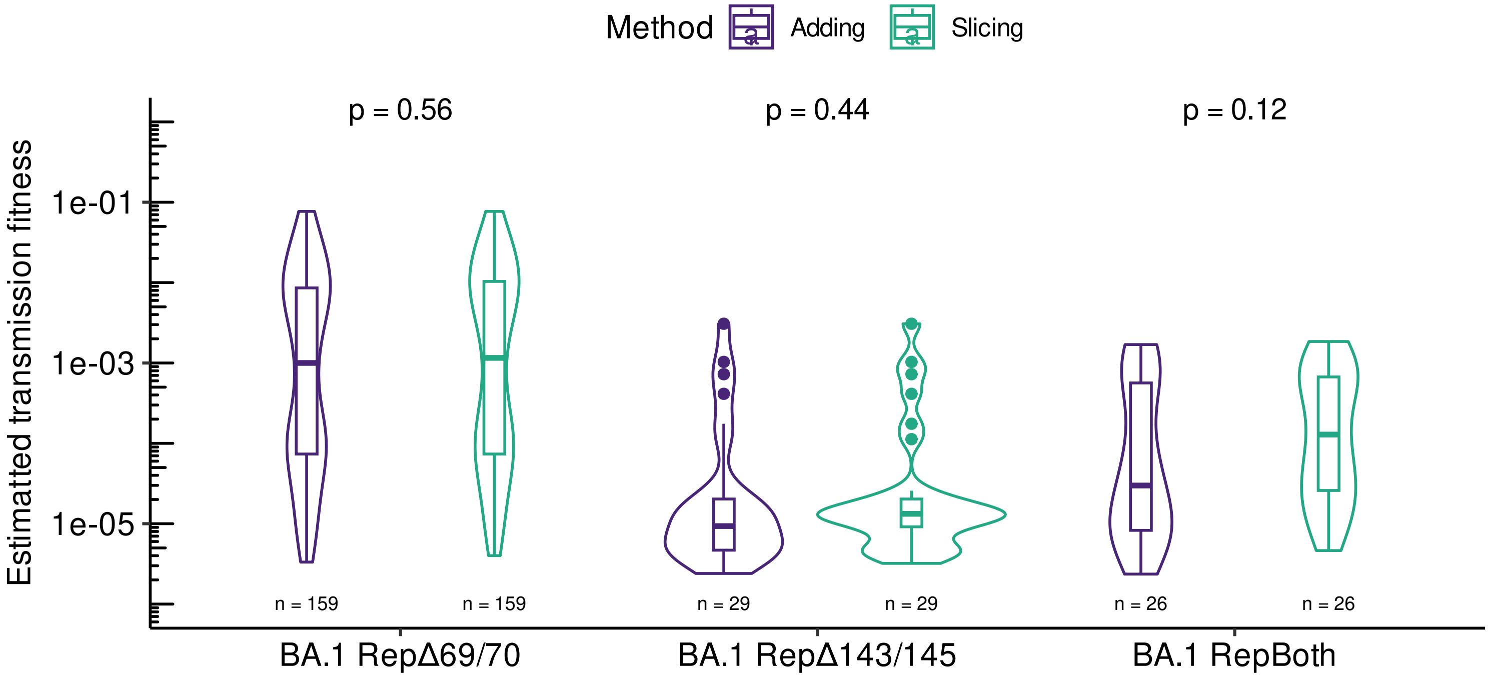


Supplemental Figure 1. Comparison of settings for estimating transmission fitness based on the transmitted cases per GISAID submission in the same time window and country of sampling, with a symmetrical time window padding of 7 days, for deletion repair haplotypes in BA.1 background. Differences considering the subdivision of time windows into country-specific sub-windows ("slicing" and "adding") are reported. P-values were calculated with the Wilcoxon rank-sum test.
