## Supplemental Figure 2 for "Genome data artifacts and functional studies of deletion repair in the BA.1 SARS-CoV-2 spike protein"

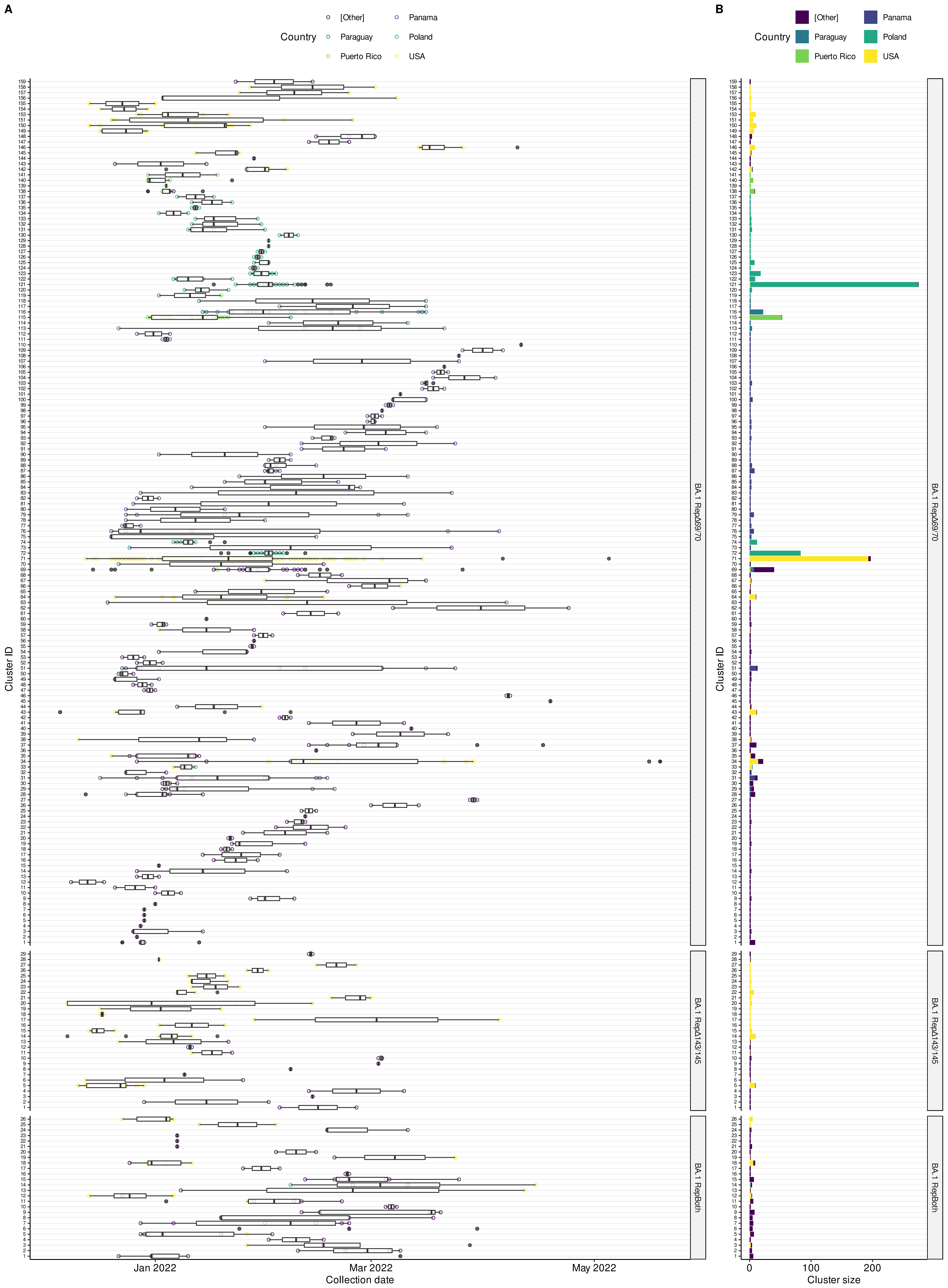


Supplemental Figure 2. Collection date and country of sampling of transmitted samples assigned to deletion repair haplotypes in BA.1 background from the GISAID survey.
