## Supplemental Figure 3 for "Genome data artifacts and functional studies of deletion repair in the BA.1 SARS-CoV-2 spike protein"

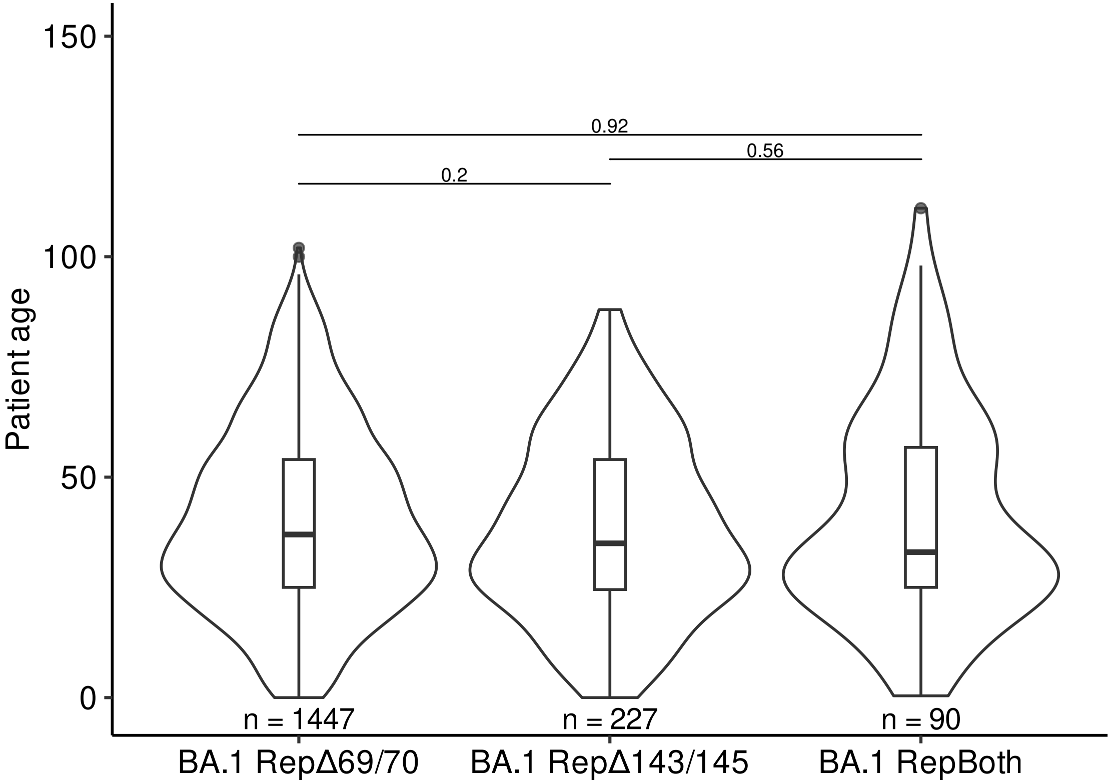


Supplemental Figure 3. Differences in patient age between deletion repair haplotypes BA.1 background from the GISAID survey. P-values of pairwise differences were calculated with the Wilcoxon rank-sum test.
