## Supplemental Figure 4 for "Genome data artifacts and functional studies of deletion repair in the BA.1 SARS-CoV-2 spike protein"

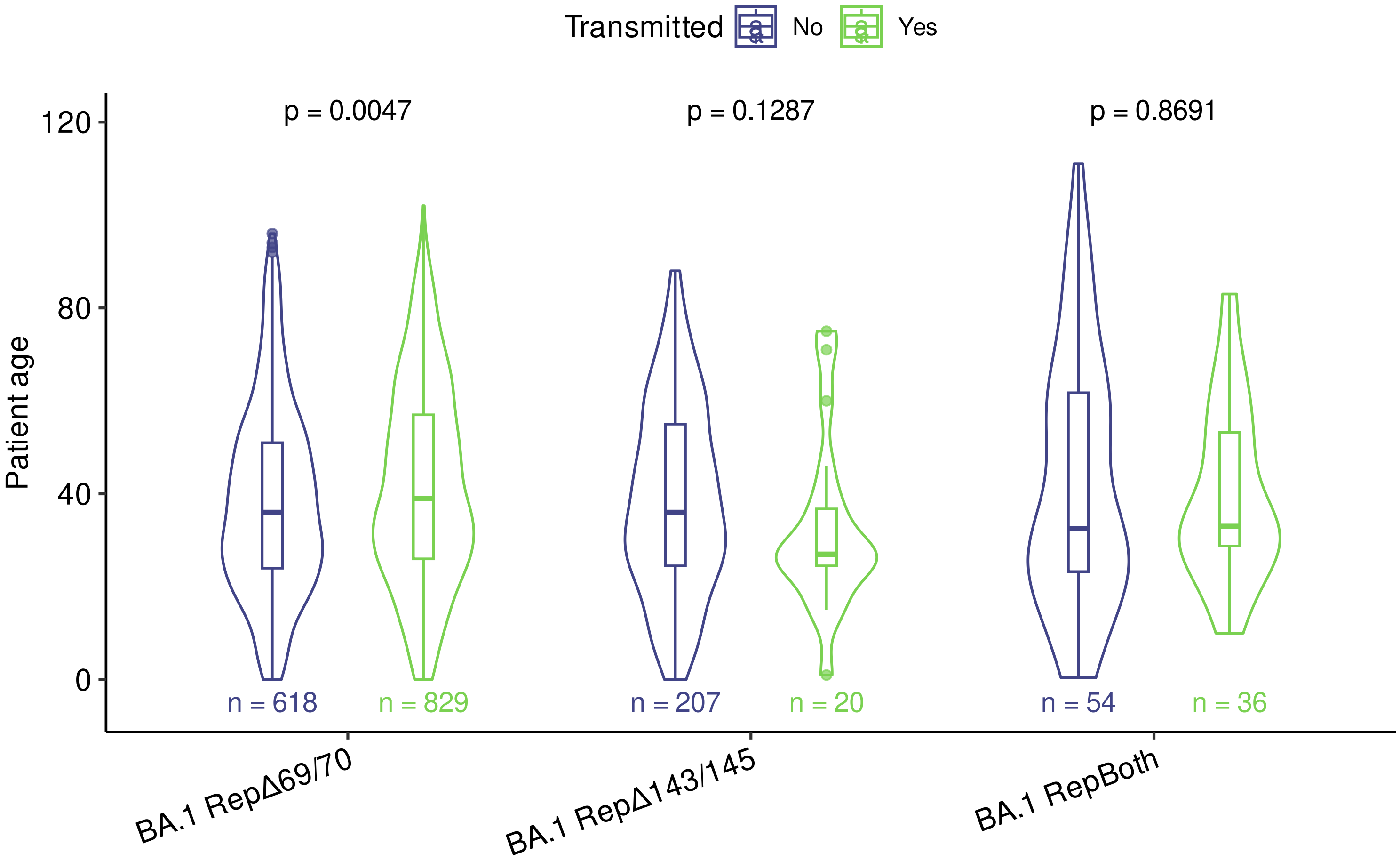


Supplemental Figure 4. Differences in within-haplotype host age, grouping samples by their involvementin transmission clusters, of deletion repair haplotypes in BA.1 background from the GISAID survey. Each data point corresponds to a different sample with known host age. P-values were calculated with the Wilcoxon rank-sum test.
