## Supplemental Figure 5 for "Genome data artifacts and functional studies of deletion repair in the BA.1 SARS-CoV-2 spike protein"

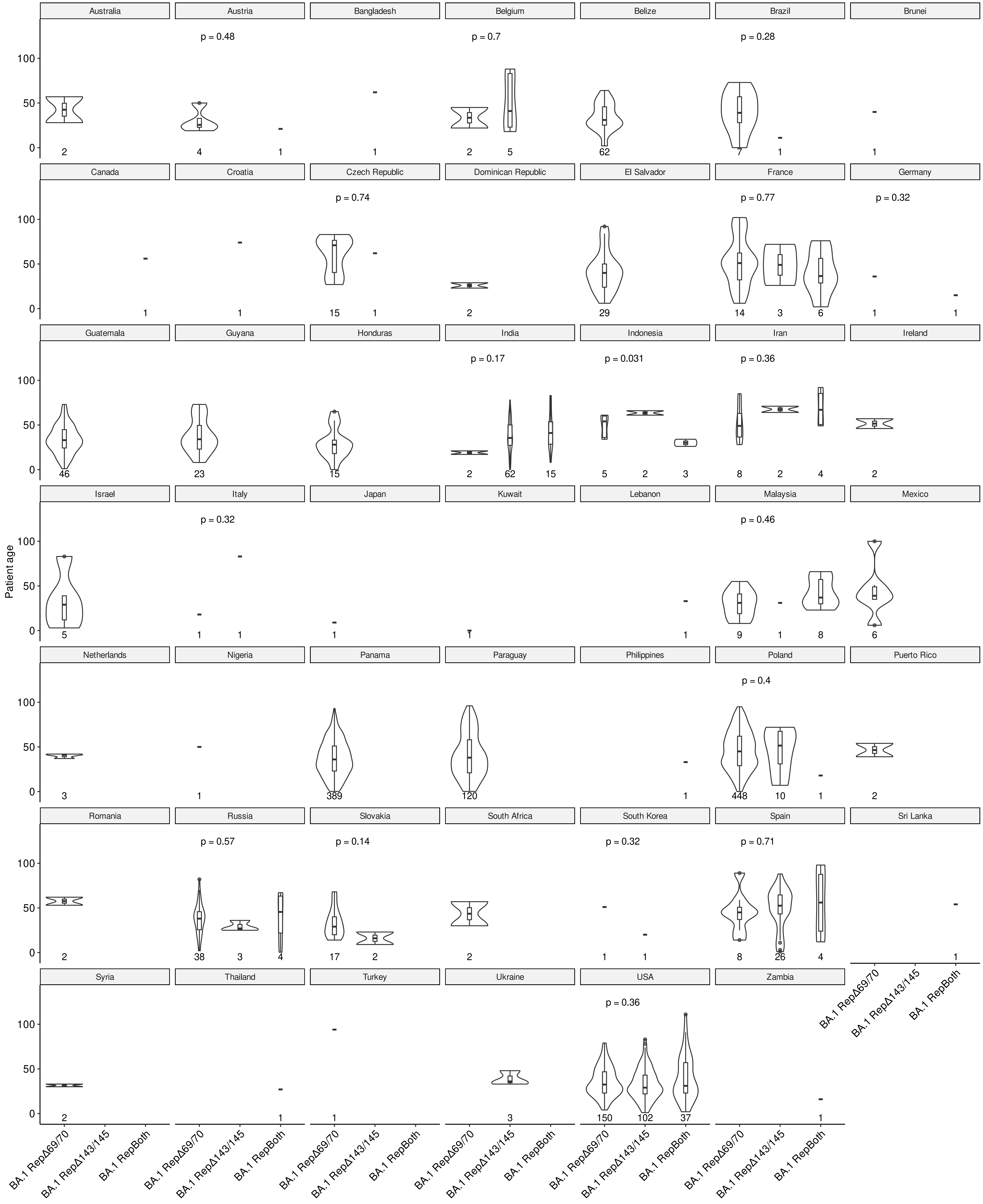

Supplemental Figure 5. Differences in patient age between deletion repair haplotypes in BA.1 background from the GISAID survey, split by country of sampling. P-values were calculated with the Wilcoxon rank-sum test.
