## Supplemental Figure 6 for "Genome data artifacts and functional studies of deletion repair in the BA.1 SARS-CoV-2 spike protein"

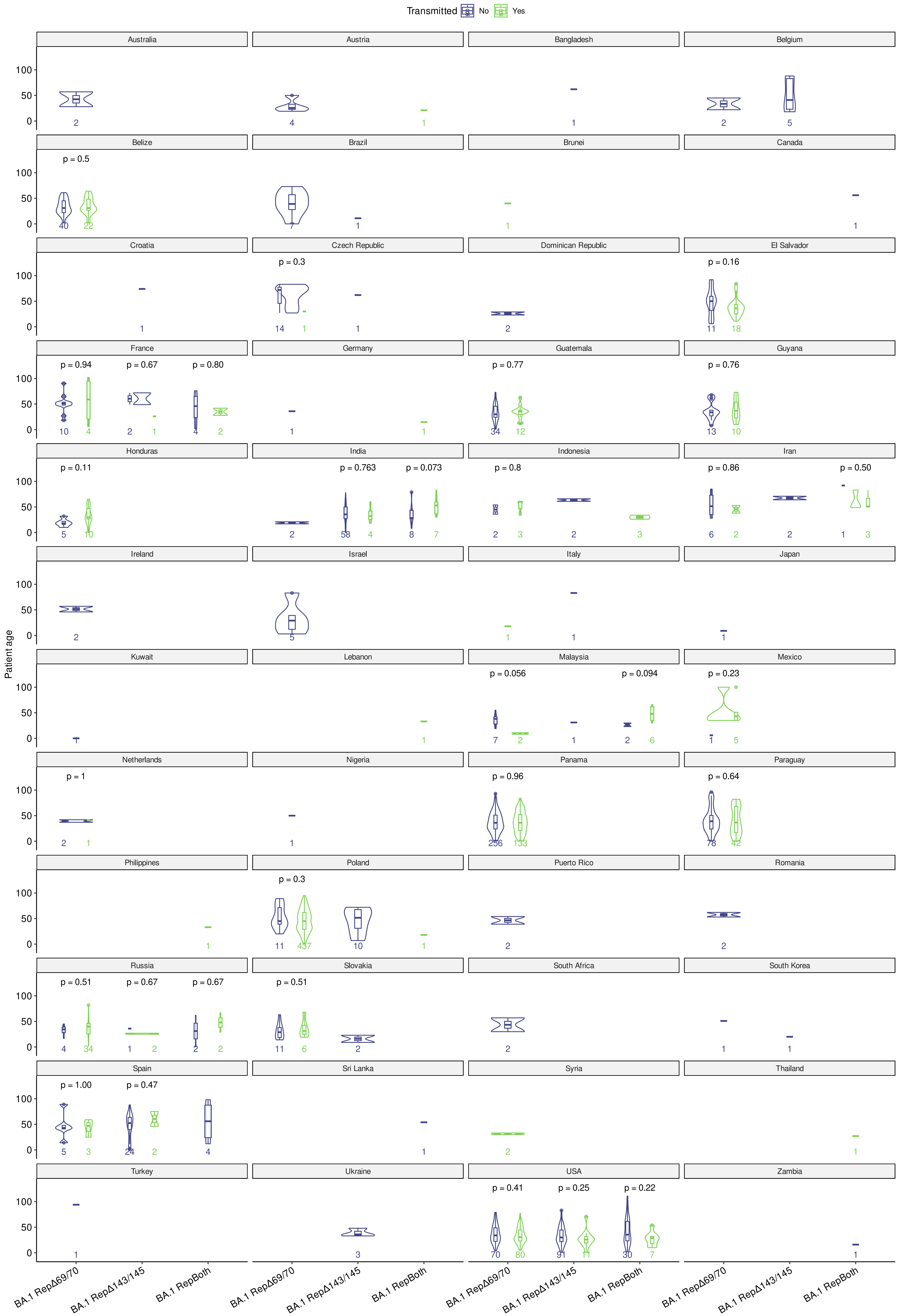


Supplemental Figure 6. Differences in within-haplotype host age, grouping samples by their involvement in transmission clusters, of deletion repair haplotypes in BA.1 background from the GISAID survey, split by country of sampling. Each data point corresponds to a different sample with known host age. P-values were calculated with the Wilcoxon rank-sum test.
