## Supplemental Figure 7 for "Genome data artifacts and functional studies of deletion repair in the BA.1 SARS-CoV-2 spike protein"

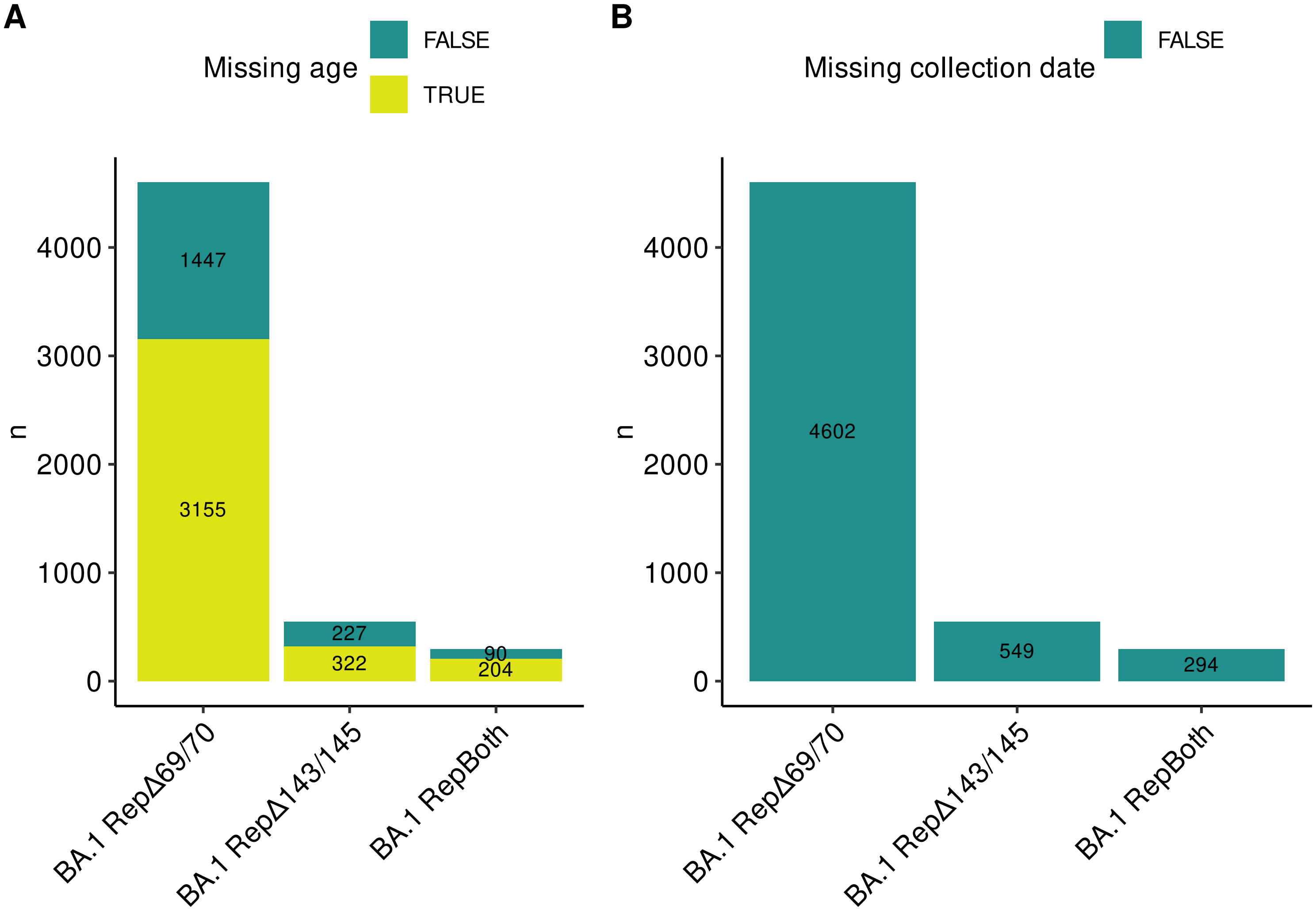


Supplemental Figure 7. Number of GISAID records (n) with missing annotation of host age (A) and collection date (B), for deletion repair haplotypes in BA.1 background.
